## Supplemental Materials for "Structural Insights into the Venus flytrap Mechanosensitive Ion Channel Flycatcher1"

**Supplemental figures:**

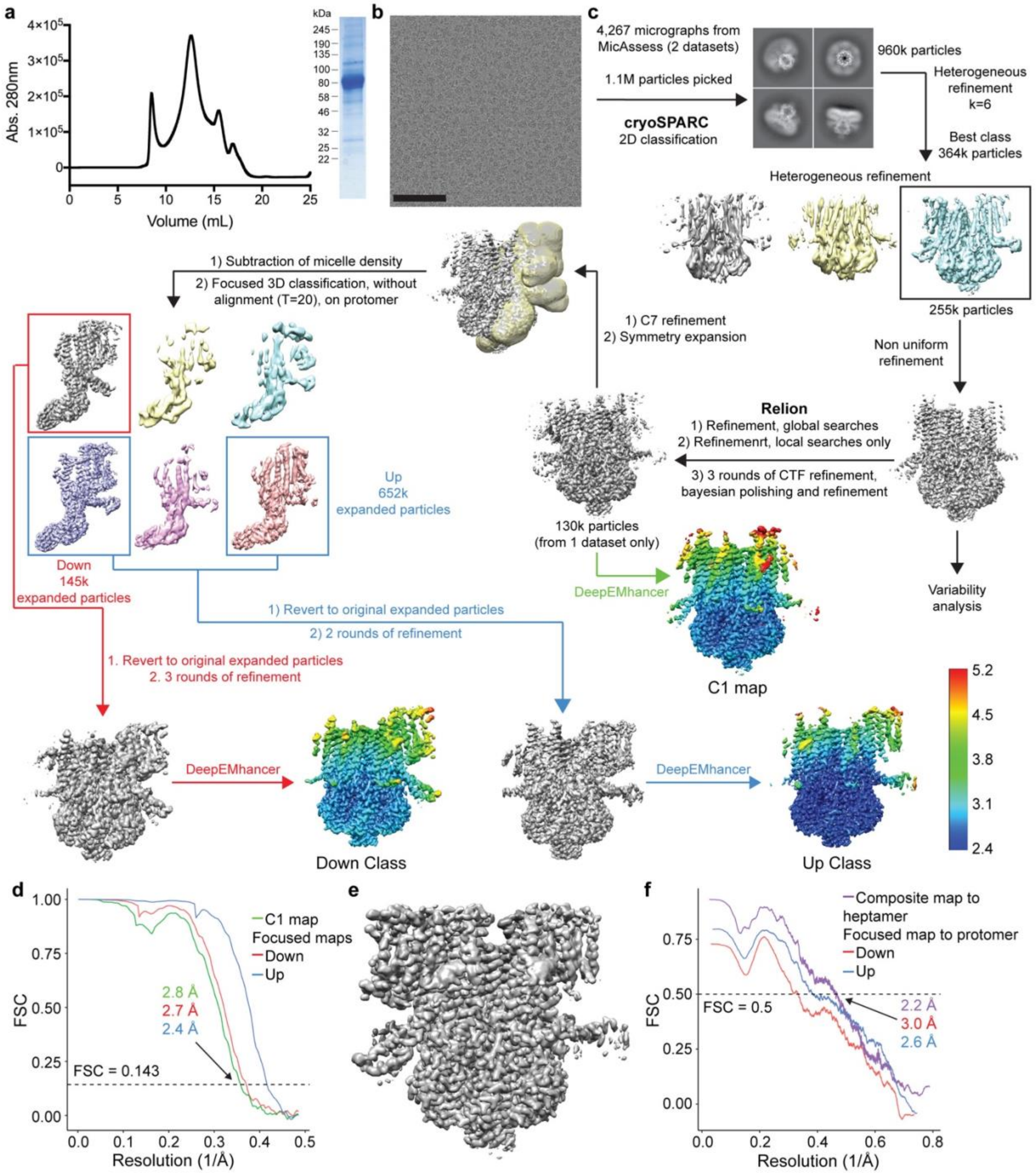


**Supplementary Figure 1: Purification and cryo-EM data processing of FLYC1.**
**a,** Size exclusion chromatography trace (left) and SDS-PAGE (right) from FLYC1 purification. Molecular weight of FLYC1 protomer is approximately 86kDa. **b,** Representative cryo-EM micrograph. Black bar is 100nm. **c,** Cryo-EM processing workflow. **d,** FSC plot of C1 and focused maps. **e,** Final composite map obtained by using the C1 and focused maps. **f,** Map to

model FSC plot of composite and focused maps. Only the protomer was used for the calculation for the focused maps.

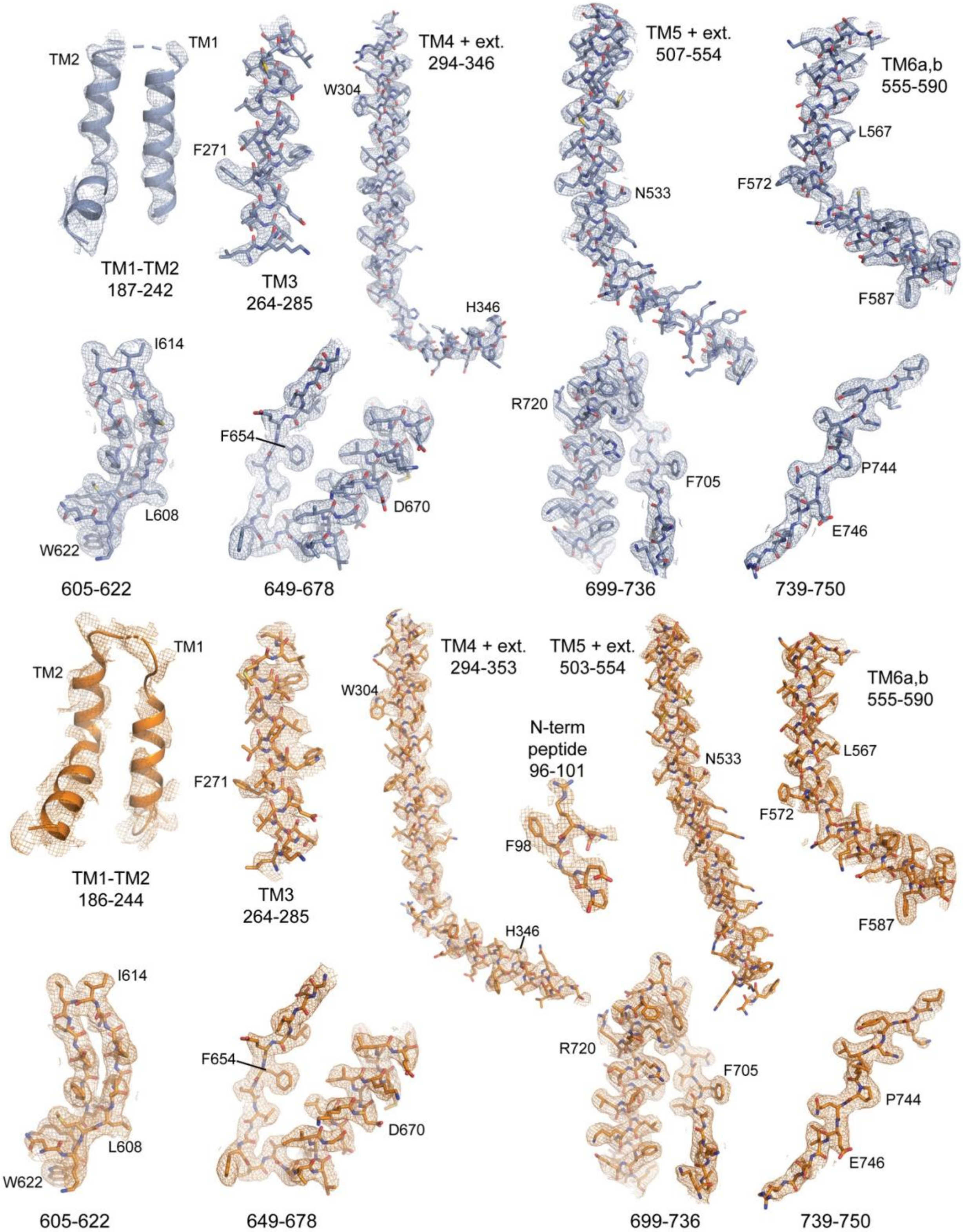


**Supplementary Figure 2: Fit of FLYC1 model to map.**
Fit of FLYC1 model for the up (top, blue) and down (bottom, orange) classes to composite map. Map was contoured at a threshold of 1 (for N-term peptide in down class) or 4 σ.

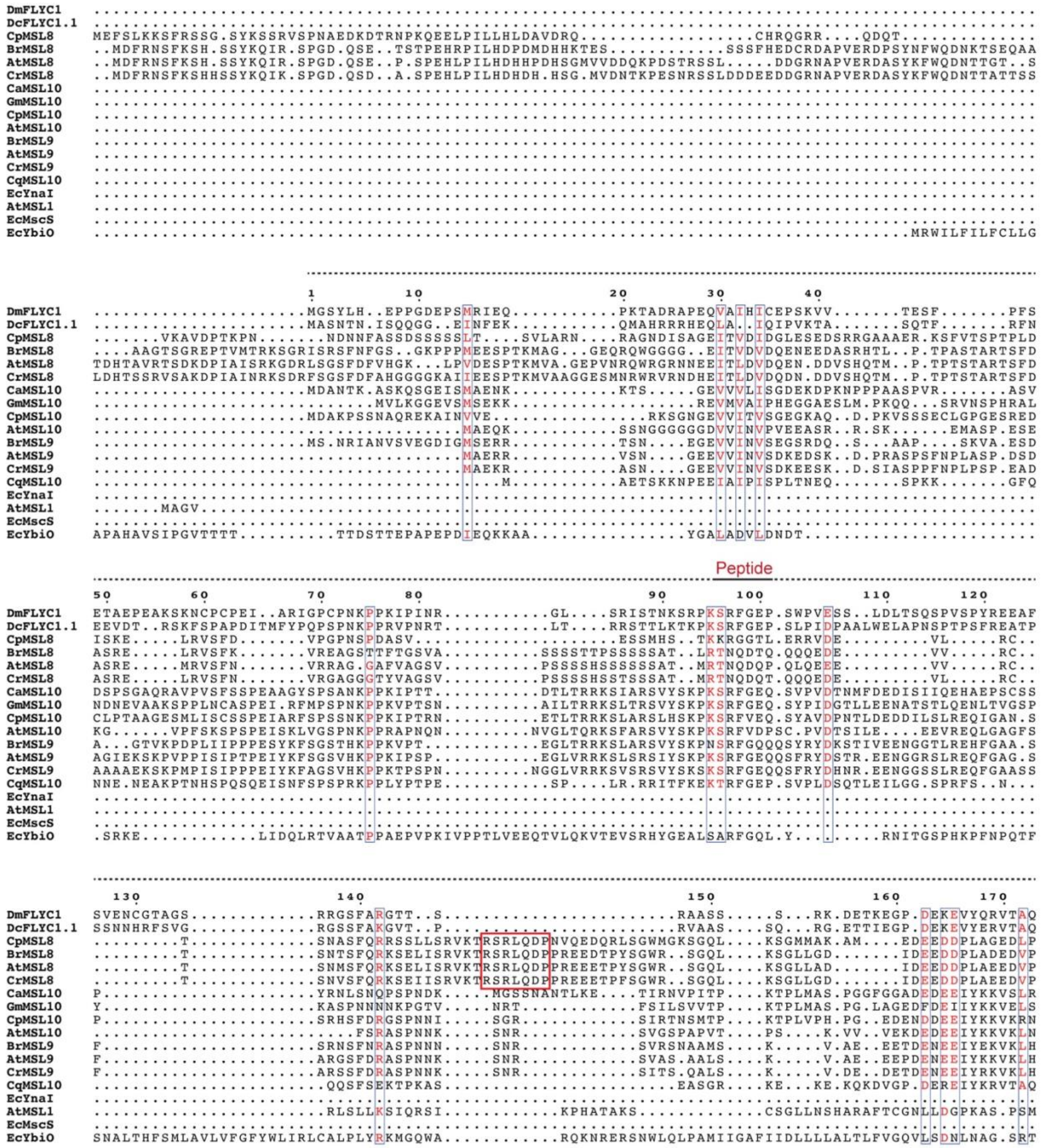
DmFLYC1
DcFLYC1.1
CpMSL8
BrMSL8
AtMSL8
CrMSL8
CaMSL10
GmMSL10
CpMSL10
AtMSL10
BrMSL9
AtMSL9
CrMSL9
CqMSL10
EcYnaI
AtMSL1
EcMscS
EcYbiO
Peptide

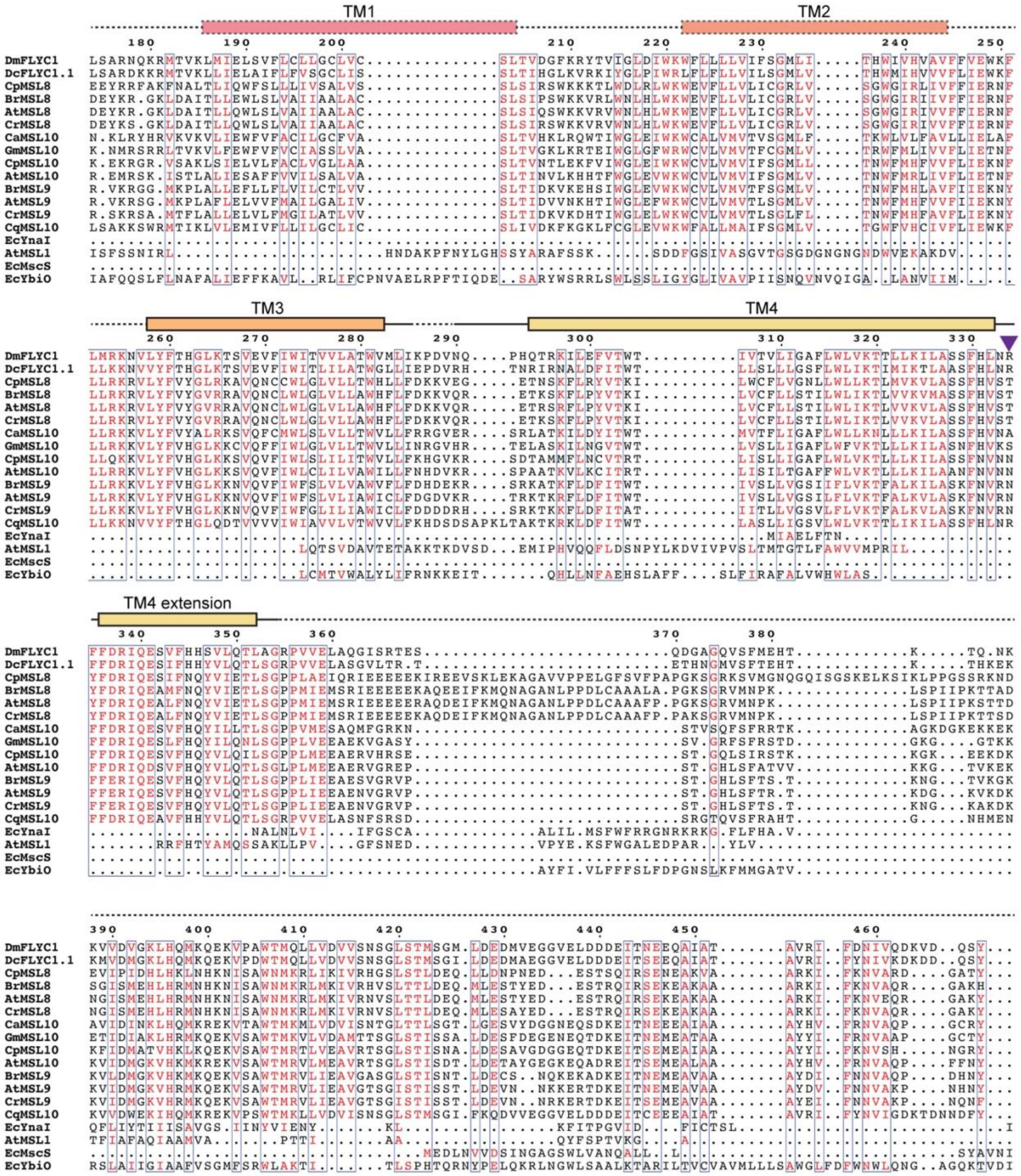
TM1
TM2
TM3
TM4
TM4 extension
DmFLYC1
DcFLYC1.1
CpMSL8
BrMSL8
AtMSL8
CrMSL8
CaMSL10
GmMSL10
CpMSL10
AtMSL10
BrMSL9
AtMSL9
CrMSL9
CqMSL10
EcYnaI
AtMSL1
EcMscS
EcYbiO

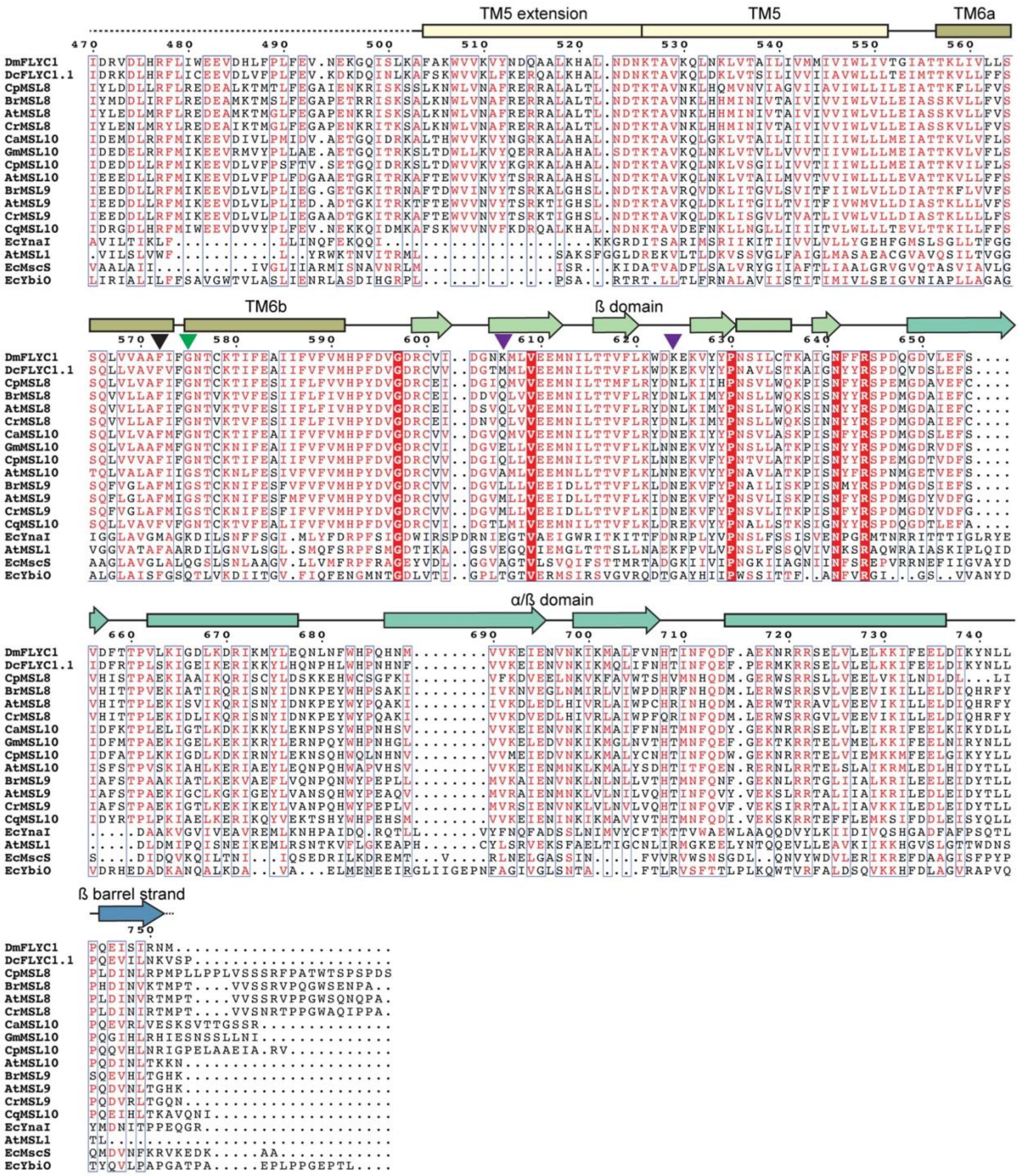


**Supplementary Figure 3. Protein sequence alignment of (Dm)FLYC1.**

Sequence alignment of DmFLYC1 (GenBank ID: QNN26181), *Drosera capensis* FLYC1.1 (QNN26182), *Carica papaya* (Cp)MSL8 (XP_021903761), *Brassica rapa* (Br)MSL8 (XP_018511053), AtMSL8 (NP_001318236), *Capsella rubella* (Cr)MSL8 isoform X1 (XP_023642637), *Coffea arabica* (Ca)MSL10-like (XP_027081862), *Glycine max* (Gm)MSL10 (XP_006578090), CpMSL10 (XP_021890723), AtMSL10 (NP_001119212), BrMSL9 (XP_009131740), AtMSL9 (NP_001331595), CrMSL9 (XP_006287126), *Chenopodium quinoa*

(Cq)MSL10-like (XP_021738929), EcYnaI (WP_000559900), AtMSL1 (NP_567165), EcMscS (WP_000389818), EcYbiO (WP_001267253). Secondary structure of FLYC1 represented at the top of each block, colored accordingly to Figure 1d. Dashed lines represent areas not built in the model. TM1 and TM2 are assigned based on TOPCONS predictions. TM4 and TM5 extensions for the TM4-5 linker. Arrow heads point to residue lining the smaller pore constriction (black), glycine hinge (green) or residues mutated in this study (purple). Red box surrounds residues of MSL8 homologs that align to the N-terminus peptide in alignment (not shown) that does not include AtMSL1 or bacterial homologs.

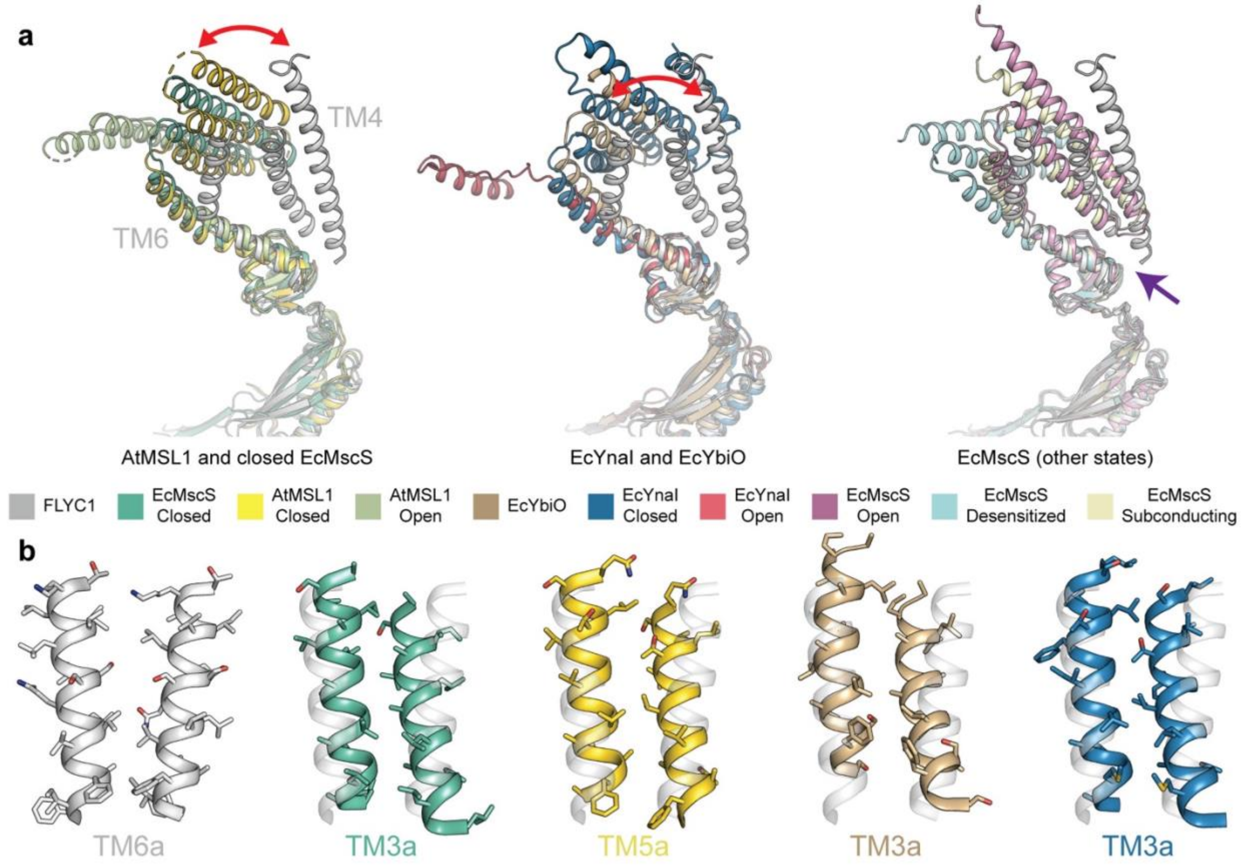


**Supplementary Figure 4. Comparison of FLYC1 with solved homologs.**
**a,** Superposition of FLYC1 subunit and reported structures of homologs (aligned on cytoplasmic domain). Red arrow show difference in rotation between FLYC1 and other homologs. Purple arrow points to the interaction between TM4-5, TM1-2 in EcMscS, and the pore helix for FLYC1 and EcMscS in both open and subconducting states. TM1-TM3 of FLYC1 are not shown. **b,** Alignment based on TM6a of two adjacent pore helices of FLYC1 and other homologs showing the presence of bulkier PDB ID) are: EcMscS closed state (2OAU), AtMSL1 closed state (6VXM), AtMSL1 open state (6VXN), EcYbiO (7A46), EcYnaI closed state (6ZYD), EcYnaI open state (6ZYE), EcMscS open state (5AJI), EcMscS desensitized state (6VYM), and EcMscS subconducting state (6VYL).

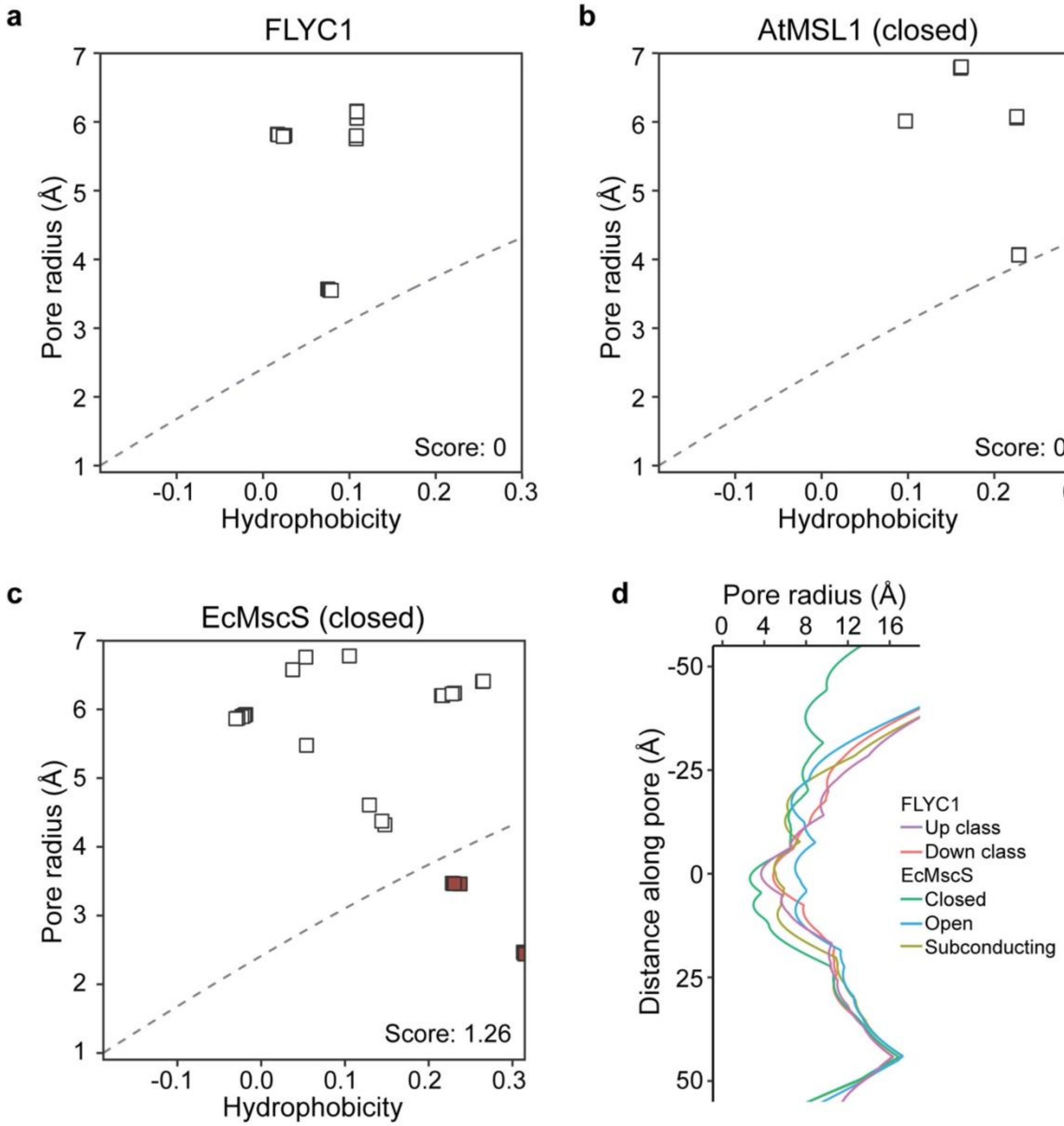


**Supplementary Figure 5. Comparison of FLYC1 and homologs pore profile.**
Heuristic prediction of likelihood of pore wetting of FLYC1 (**a**), closed AtMSL1 (**b**) and closed EcMscS (**c**). A score greater than 0.55 predicts the presence of at least one energetic barrier to water permeation in the channel pore. **d,** Pore profile of hypothetical C7 FLYC1 conformations in comparison to EcMscS states.

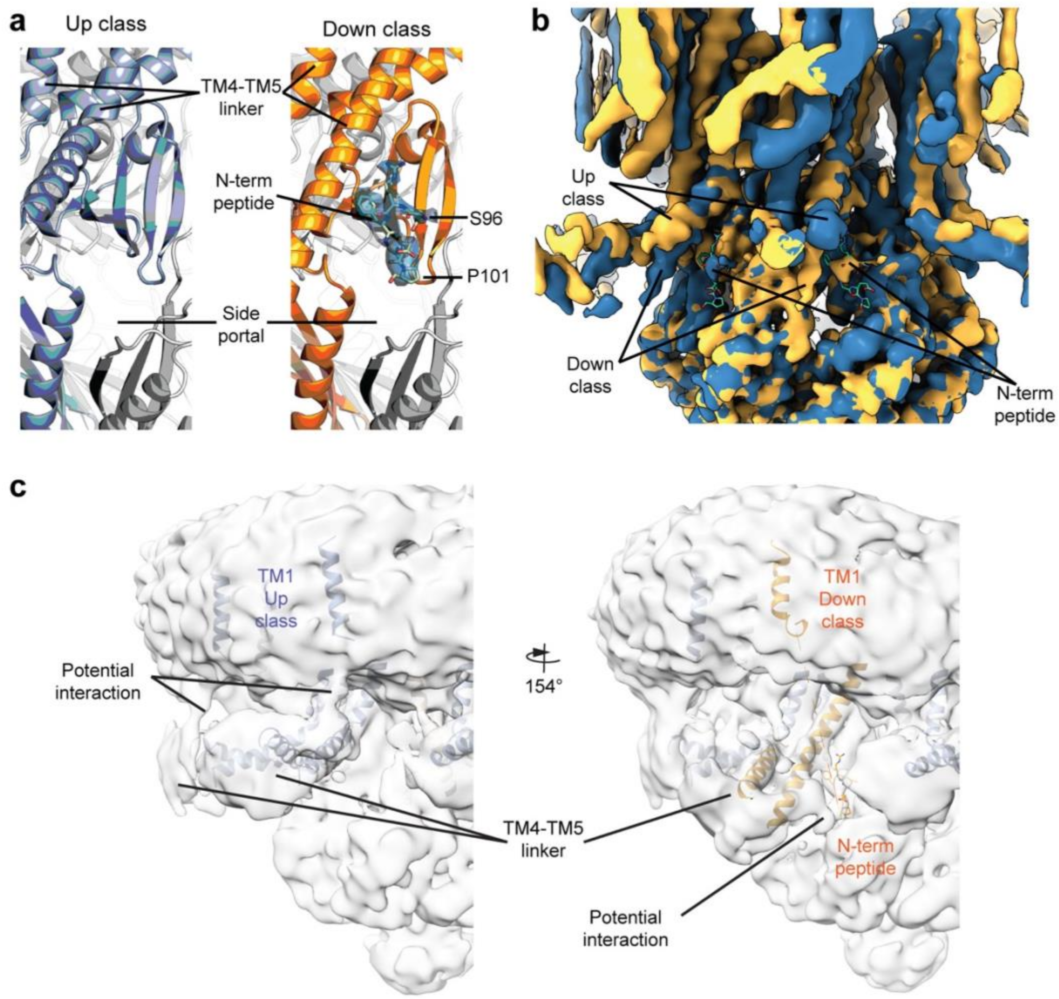


**Supplementary Figure 6. Interaction of N-terminus of FLYC1.**
**a,** Superposed first (blue) and last frame (yellow) of variability analysis volume series showing peptide density for part of the N-terminus limited to the down class. First and last residues built for the peptide are labeled. **b,** Peptide density from final reconstruction. **c,** 6Å low-pass filter unsharpened C1 map showing density for potential interaction between TM4-TM5 linker and TM1, in the up class (left), or N-terminus peptide, in the down class (right).

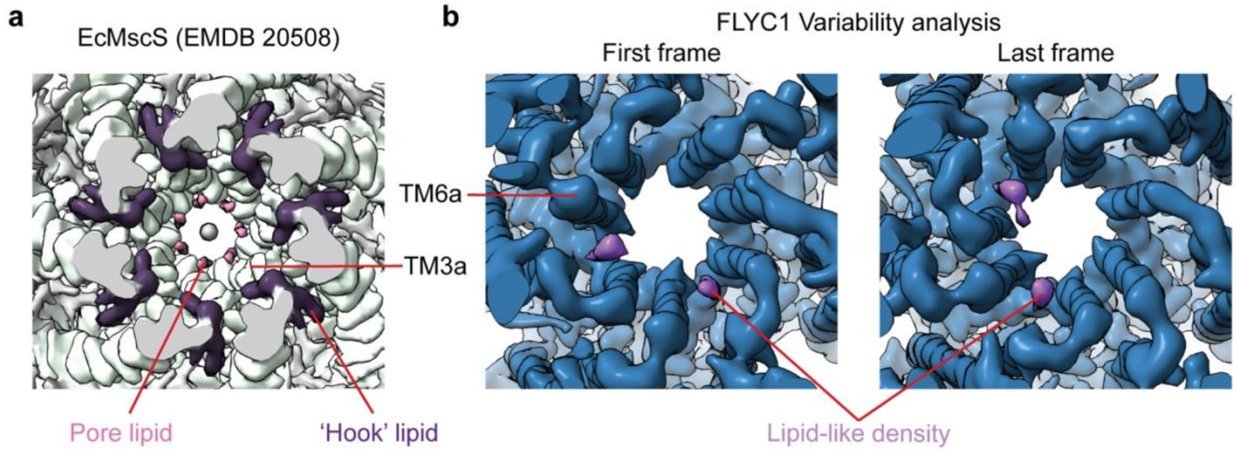


**Supplementary Figure 7. FLYC1 and EcMscS lipid-like densities at the outer face of the pore.**
**a,** Densities assigned to lipids in EcMscS map (EMDB 20508). Pore and 'hook' lipids are colored pink and dark purple, respectively. **b,** Maps corresponding to first (left) and last (right) frames of the first component from variability analysis of FLYC1. Lipid-like densities associated to protomers in the down conformation are colored in purple.

**Video 1: 3D EM Variability analysis of FLYC1**
First variability component of FLYC1 after C7 symmetry expansion. Densities have been low-pass filtered to 6Å and high-pass filtered to 20Å. Appearance of N-terminus peptide and lipid-like density at the top of the channel are shown associated to the down class.

**Supplementary Table 1. Data collection, processing, model refinement and validation.**

| | C1 map | Down focused map | Up focused map | Composite map |
|---|---|---|---|---|
| **Data collection and processing** | | | | |
| Magnification | 29000 | | | |
| Voltage (kV) | 300 | | | |
| Electron exposure (e–/Å$^2$) | 50 | | | |
| Defocus range (μm) | -0.7 to -2.2 | | | |
| Pixel size (Å) | 1.03 | | | |
| Initial particle images (no.) | 959,656 | | | |
| Symmetry imposed | C1 | C1 | C1 | C1 |
| Final particle images (no.) | 129,933 | 145,362[*] | 651,815[*] | |
| Map resolution (Å) | 2.8 | 2.7 | 2.4 | 2.8[†] |
| FSC threshold | 0.143 | 0.143 | 0.143 | 0.143 |
| Map sharpening *B* factor (Å$^2$) | -47 | -43 | -38 | |
| **Model** | | | | |
| Composition | | | | |
| Peptide chains | 7 | 1[‡] | 1[‡] | 7 |
| Protein residues | 2176 | 386 | 371 | 2612 |
| Ligands | 7 | 1 | 1 | 7 |
| R.m.s. deviations | | | | |
| Bond lengths (Å) | 0.023 | 0.022 | 0.022 | 0.022 |
| Bond angles (°) | 1.576 | 1.534 | 1.531 | 1.536 |
| Validation | | | | |
| MolProbity score | 0.59 | 0.56 | 0.57 | 0.58 |
| Clashscore | 0.25 | 0.16 | 0.17 | 0.22 |
| EMRinger score | 4.07 | 3.32 | 3.64 | 4.14 |
| Poor rotamers (%) | 0.00 | 0.00 | 0.00 | 0.00 |
| Ramachandran plot | | | | |
| Favored (%) | 98.84 | 98.66 | 99.17 | 98.94 |
| Allowed (%) | 1.16 | 1.34 | 0.83 | 1.06 |
| Disallowed (%) | 0.00 | 0.00 | 0.00 | 0.00 |
| **Deposition ID** | | | | |
| EMDB | 24187 | 24188 | 24189 | 24186 |
| PDB | 7N5E | 7N5F | 7N5G | 7N5D |

[*] After symmetry expansion.
[†] Assigned resolution. Due to the final map being a composite map, resolution estimation based on FSC between half-maps could not be calculated.
[‡] Only protomer was used for model validation in the focused maps.
